# Development of a targeted sequencing approach to identify prognostic, predictive and diagnostic markers in paediatric solid tumours

**DOI:** 10.1101/165746

**Authors:** Elisa Izquierdo, Lina Yuan, Sally George, Michael Hubank, Chris Jones, Paula Proszek, Janet Shipley, Susanne A Gatz, Caedyn Stinson, Andrew S. Moore, Steven C. Clifford, Debbie Hicks, Janet Lindsey, Rebecca Hill, Thomas S. Jacques, Jane Chalker, Khin Thway, Simon O’Connor, Lynley Marshall, Lucas Moreno, Andrew Pearson, Louis Chesler, Brian A. Walker, David Gonzalez De Castro

## Abstract

The implementation of personalised medicine in childhood cancers has been limited by a lack of clinically validated multi-target sequencing approaches specific for paediatric solid tumours. In order to support innovative clinical trials in high-risk patients with unmet need, we have developed a clinically relevant targeted sequencing panel spanning 311 kb and comprising 78 genes involved in childhood cancers. A total of 132 samples were used for the validation of the panel, including Horizon Discovery cell blends (n=4), cell lines (n=15), formalin-fixed paraffin embedded (FFPE, n=83) and fresh frozen tissue (FF, n=30) patient samples. Cell blends containing known single nucleotide variants (SNVs, n=528) and small insertion-deletions (indels n=108) were used to define panel sensitivities of ≥98% for SNVs and ≥83% for indels [95% CI] and panel specificity of ≥98% [95% CI] for SNVs. FFPE samples performed comparably to FF samples (n=15 paired). Of 95 well-characterised genetic abnormalities in 33 clinical specimens and 13 cell lines (including SNVs, indels, amplifications, rearrangements and chromosome losses), 94 (98.9%) were detected by our approach. We have validated a robust and practical methodology to guide clinical management of children with solid tumours based on their molecular profiles. Our work demonstrates the value of targeted gene sequencing in the development of precision medicine strategies in paediatric oncology.

## Introduction

Cancer remains the leading cause of death due to disease in children aged >1 year [1]. Cure rates for paediatric solid tumours have not substantially improved in the past decade with patients having recurrent disease performing particularly badly, reflecting the limitations of current approaches that employ intensive chemotherapy, surgery and radiation [2–4]. In adults, the stratification of patients by genetic profiling using high throughput sequencing has supported adaptive clinical trials [5, 6], and there is an urgent need to translate such opportunities to the treatment of childhood disease.

The genomic landscape of paediatric cancer is becoming increasingly well-defined leading to the conclusion that childhood cancers have in general fewer somatic mutations than adults, but that mutations in epigenetic regulators occur at a higher incidence [7–17]. Key recent findings include recurrent mutations in the genes encoding histones 3.3 and 3.1 (*H3F3A* and *HIST1H3B*) as well as the activin A receptor type I (*ACVR1*) that are unique to paediatric high-grade glioma (pHGG) and diffuse intrinsic pontine glioma (DIPG) [18–20]. Similarly, *ATRX* mutations, *TERT* rearrangements and *MYCN* amplification define mutually exclusive molecular subgroups of neuroblastoma, all of which are associated with poor prognosis [21–23]. The newly proposed molecular-based medulloblastoma sub-classification defines subgroups, each of which potentially requires a tailored therapeutic strategy [7, 11, 24].

Despite our improved knowledge of somatic alterations in paediatric cancers, precision medicine remains unavailable for the majority of patients. For example, a small number of early-phase paediatric trials are recruiting children whose tumours harbour genetic alterations including *ALK* genomic alterations (mutations, amplifications or translocations) that can be treated with ALK inhibitors and *BRAF* V600 mutant tumours that can be treated with BRAF or MEK inhibitors.

In addition, there is now an extensive list of recurrent genetic alterations with potential diagnostic, prognostic or predictive value, and sequential testing of single genes using standard methods has become unfeasible due to lack of available material and high costs. High-throughput sequencing (also known as next generation sequencing or NGS) offers a solution to these issues. In particular, panel-based NGS assays which simultaneously sequence a targeted set of genes with recurrent alterations, associated with known clinical or biological implications are cheaper, less challenging in terms of interpretation and more suited to clinical diagnostics than current approaches [25]. Despite this, development and validation of high throughput gene panel sequencing is challenging. Typically, DNA is only available from formalin-fixed, paraffin-embedded (FFPE) samples, which yields relatively poor quality DNA. DNA extraction and library construction to clinical laboratory standards requires optimisation, and it is necessary to construct a standardised informatics pipeline that identifies and interprets actionable mutations. Appropriate and rapid clinical reporting of identified variants and incorporation of the results into the electronic patient records also need to be considered if molecular stratification of childhood cancer is to be successfully translated to the clinic [26]. There are several examples of validation and implementation of targeted sequencing in adult cancer [27–30]. In the past two years, several approaches using high-throughput sequencing have been applied for clinical decision-making in children with solid tumours [31–34], however a clinically validated panel specifically targeting recurrent alterations in childhood cancers using archival FFPE specimens would significantly assist the development of molecular stratification strategies in paediatric oncology.

Here we describe the development and validation, within an accredited clinical pathology laboratory (CPA UK), of a paediatric solid tumour sequencing panel for use with either routine FFPE or fresh frozen (FF) samples. As part of the validation, we established overall performance, sensitivity, specificity, repeatability, reproducibility, accuracy and limit of detection, following guidelines previously described for validation of genetic tests [35].

## Results

### Selection of Panel Content

The panel design covers a total of 78 genes (Table 1), either recurrently altered in paediatric cancers or clinically actionable in adult cancers and with potential application in childhood tumours. The genes were selected in wide-collaboration with national experts in paediatric oncology patient care covering all areas of paediatric solid tumours (glioma, medulloblastoma, bone sarcomas, soft tissue sarcomas, renal tumours and neuroblastoma among others). Targets were chosen by consensus based on most clinically relevant aberrations. Factors influencing the choice of targets included: childhood tumour type where alterations have been reported, molecules targeting these genes and clinical trials available for children with solid tumours (Supplemental Table S1 A). A library of customized biotinylated DNA probes was designed to capture a total of ~311(kilobase) kb for the detection of single nucleotide variants (SNVs), short insertion-deletions (indels), copy number variations and structural rearrangements (Supplemental Table S1 B). Exons were padded with 5 base pairs (bp) of intronic sequence to increase exon depth and for detection of splice-site variants.

**Table 1.** Gene panel list including 78 genes recurrently altered in paediatric cancers or clinically actionable.

|  |  |  |  |
| --- | --- | --- | --- |
| *ACVR1* | *CTNNB1* | *IL3* | *PPM1D* |
| *AKT1* | *DDR2* | *IL6* | *PTCH1* |
| *ALK* | *DDX3X* | *KIT* | *PTEN* |
| *AMER1* | *DICER1* | *KMT2D* | *PTPN11* |
| *APC* | *EGFR* | *KRAS* | *PTPRD* |
| *ARID1A* | *ERBB2* | *LMO1* | *RB1* |
| *ARID1B* | *ERG* | *MAP2K1* | *RET* |
| *ASXL1* | *ETV6* | *MAP2K2* | *ROS1* |
| *ATM* | *EWSR1* | *MDM2* | *SETD2* |
| *ATRX* | *FBXW7* | *MYCN* | *SMARCA4* |
| *BARD1* | *FGFR1* | *MYOD1* | *SMARCB1* |
| *BCOR* | *FGFR2* | *NCOA2* | *SS18* |
| *BRAF* | *FGFR3* | *NF1* | *SUFU* |
| *CASC15* | *FGFR4* | *NRAS* | *TENM3* |
| *CDK4* | *FUS* | *PAX3* | *TP53* |
| *CDK6* | *H3F3A* | *PAX7* | *TSC1* |
| *CDKN2A* | *HIST1H3B* | *PDGFRA* | *WT1* |
| *CDKN2B* | *HIST1H3C* | *PHOX2B* | *ZHX2* |
| *CFL1* | *HRAS* | *PIK3CA* |  |
| *CHEK2* | *IDH1* | *PIK3R1* |  |

### Panel Validation

Research use of sequence capture assays has become common, but basing clinical care on gene panel sequencing results requires confident calling of both variant and non-variant sequence, and a full understanding of the performance of the assay. Implementation in the clinic therefore requires robust validation in an accredited laboratory.

To validate the paediatric gene panel, we followed the standardised framework for clinical assay validation set out by Mattocks *et al*. [35]. We determined overall performance of the panel across the target regions, measuring precision, sensitivity and specificity. As a standard, we used a set of four Horizon cell blends previously characterized by NGS and droplet digital PCR (ddPCR) (Supplemental Table S2 A and S2 B) and 15 paediatric cell lines with known variants. For further validation, we performed capture and sequencing on 83 FFPE and 30 FF clinical samples (Supplemental Table S3).

### Overall Performance

Overall, the panel performed well, with over 96% of 901 regions of interest achieving specification. Only 24 (2.7%) regions were classified as underperforming across the four cell blends and five FFPE samples, with read depth lower than 2 x standard deviation (SD) of the mean based on log_2_ (Supplemental Tables S4 A and S4 B). 22 of 24 underperforming regions were located within highly GC-enriched regions, which are known to be refractory to efficient hybridization and/or amplification (Figure 1 and Supplemental Table S4 C).

**Figure 1.**
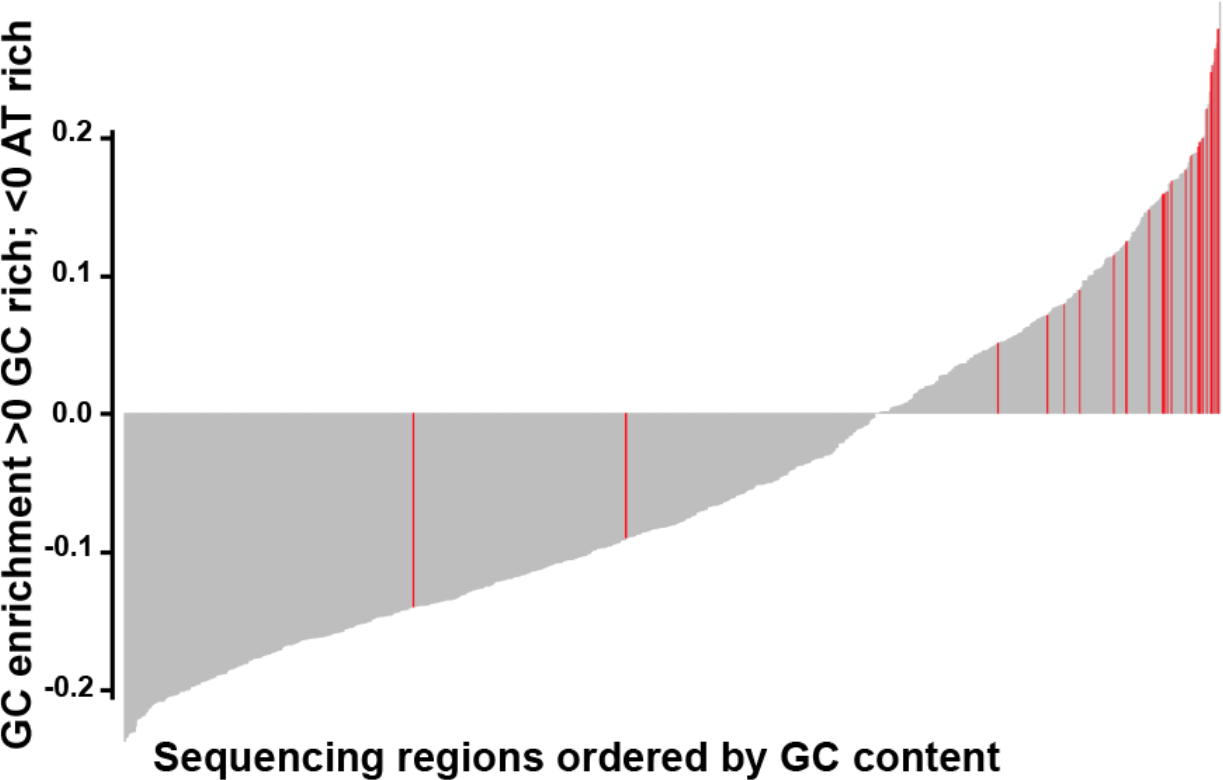
Bar plot showing GC content in the 901 regions capture by the panel. The plot is ordered from low to high GC-content of each region capture. Red bars highlight the underperforming regions (24/901), mainly located within GC-enriched regions.

Quality and coverage metrics were generated across all samples (Supplemental Table S5 and S6). The average total number of reads was 8.8x10^6^ (SD=3.1x10^6^) for FFPE and 7.9x10^6^ (SD=3x10^6^) for high molecular weight (HMW) samples (FF and cell lines). The percentage mapped (96.1±3.9 for FFPE vs 97.3±2.5 for high molecular weight samples) and percentage of bases from unique reads on target (45.9±3 for FFPE vs 42.7±2.4 for HMW) was very similar for both FFPE and HMW samples. Duplicates were higher in FFPE samples (60.2% for FFPE vs 36.1% for HMW). The overall mean depth was 698 ± 365 for FFPE vs 899 ± 347 for HMW (Table 2).

**Table 2.** Average quality metrics across all samples. Data expressed as means ± standard deviation.

|  | *Total reads* | *Percentage of reads mapped* | *Percentage of duplicates* | *Percentage of unique on target* | *Mean depth* |
| --- | --- | --- | --- | --- | --- |
| ***FFPE (n=83)*** | $8.8 \times 10^6 \pm 3.1 \times 10^6$ | 96.1±3.9 | 60.2±13.7 | 45.9±3 | 698±365 |
| ***FF and cell lines (n=49)*** | $7.9 \times 10^6 \pm 3 \times 10^6$ | 97.3±2.5 | 36.1±9.7 | 42.7±2.4 | 899±347 |

### Limit of detection

To determine the limit of detection, SNVs present in the cell blends at known variant allele frequency (VAF) were used. The pipeline detected all 61 SNVs including 33 SNVs with an expected VAF of 4-5%. 15/17 expected indels were detected. Of the two indels not detected, one was 18 bp in length at an expected VAF of 4.2%, whilst the other was 2 bp at 5% VAF (Figure 2 and Supplemental Table S7 A). We therefore established a minimum threshold of 5% VAF in the analysis pipeline, which allows for detection of a heterozygous mutation when >10% neoplastic cells are present in the tumour sample.

**Figure 2.**
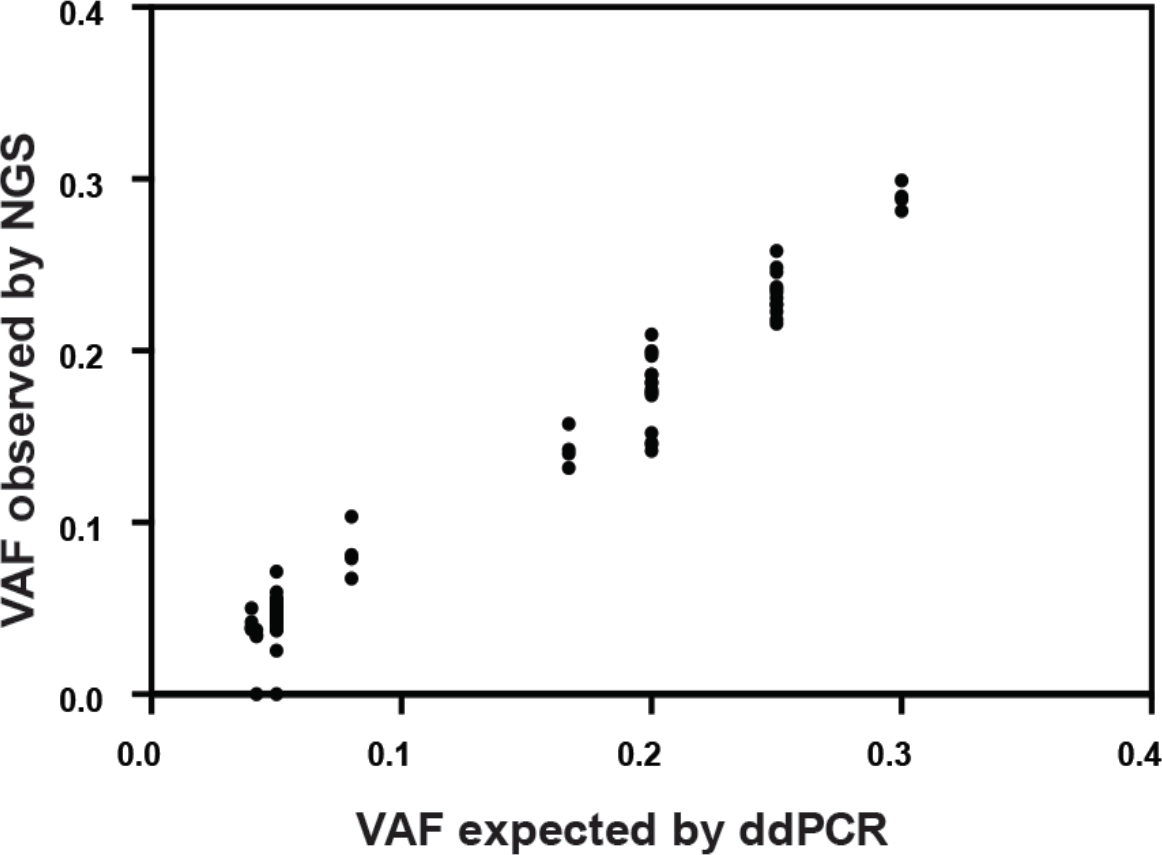
Comparison of known variant allele frequencies by droplet digital PCR (x axis) against variant allele frequency obtained by NGS (y axis) for all cancer-specific variants (61 single nucleotide variants, SNVs and 17 insertion-deletions, indels). Overall correlation was r^2^=0.969 [95% CI: 0.975-0.990; p<0.0001].

### Assessment of Precision

To measure precision, we took advantage of natural variants present as intrinsic “background” SNVs and indels in the captured regions from the four cell blends. Precision was assessed by comparing the alterations expected with those detected to obtain within run-precision (repeatability), and between run-precision data (intermediate precision). Variants ≤ 5% in all four blends and within poor performing regions were excluded leaving a total of 528 SNVs (132 variants in 4 blends) and 108 indels (27 indels in 4 blends) for analysis. All of the 528 SNVs and 90 out of 108 (83%) indels were detected (Supplemental Table S8).

#### Repeatability

Pairwise correlation of VAF between runs was r^2^≥0.994 [95%CI:0.991-0.996] for SNVs and r^2^≥0.785 [95%CI:0.652-0.919] for indels (Supplemental Figure S1 and Supplemental S2) indicating that the panel accurately reproduces data from repeat samples on the same run.

#### Intermediate precision

Pairwise correlation was r^2^≥0.995 [95%CI:0.993-0.997] and overall correlation was r^2^=0.996 [95%CI:0.995-0.997] for SNV detection. For indels pairwise correlation was r^2^≥0.827 [95%CI: 0.716-0.937] and overall correlation was r^2^≥0.875 [95%CI:0.829-0.921] (Table 3 and Supplemental Figure S3 and Supplemental Figure S4) indicating that the panel accurately reproduces data from repeat samples on different runs.

**Table 3.**
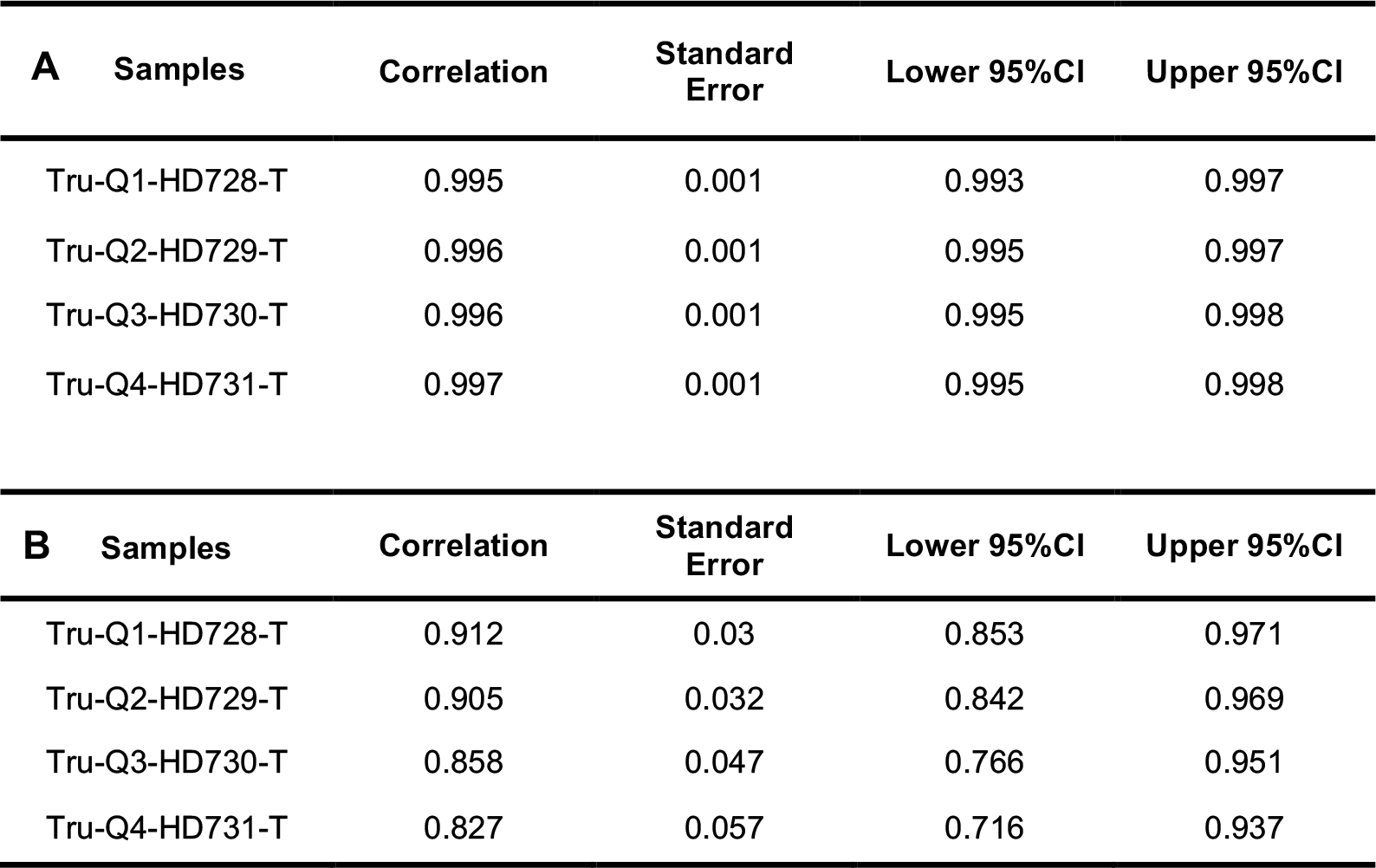
Pairwise correlation of (A) single nucleotide variants (SNVs) and (B) insertion-deletions (indels) for each of the 4 cell blends with identical background variants between the two runs.

| **A** | **Samples** | **Correlation** | **Standard Error** | **Lower 95%CI** | **Upper 95%CI** |
| --- | --- | --- | --- | --- | --- |
|  | Tru-Q1-HD728-T | 0.995 | 0.001 | 0.993 | 0.997 |
|  | Tru-Q2-HD729-T | 0.996 | 0.001 | 0.995 | 0.997 |
|  | Tru-Q3-HD730-T | 0.996 | 0.001 | 0.995 | 0.998 |
|  | Tru-Q4-HD731-T | 0.997 | 0.001 | 0.995 | 0.998 |

| **B** | **Samples** | **Correlation** | **Standard Error** | **Lower 95%CI** | **Upper 95%CI** |
| --- | --- | --- | --- | --- | --- |
|  | Tru-Q1-HD728-T | 0.912 | 0.03 | 0.853 | 0.971 |
|  | Tru-Q2-HD729-T | 0.905 | 0.032 | 0.842 | 0.969 |
|  | Tru-Q3-HD730-T | 0.858 | 0.047 | 0.766 | 0.951 |
|  | Tru-Q4-HD731-T | 0.827 | 0.057 | 0.716 | 0.937 |

### Assessment of sensitivity and specificity

To determine sensitivity we used the same background 528 SNVs and 108 indels, together with the known cancer-specific variants (61 SNVs and 17 indels) from the four cell blends. SNVs and indels were called and their presence was compared to the list of variants expected in the capture regions from the cell blends (Supplemental Table S7 A and Supplemental Table S8). All the SNVs were detected, resulting in a sensitivity of ≥98% [95%CI:0.98-1]. From the 108 background indels, 18 were not detected, as were 2 of the cancer-specific indels, obtaining a sensitivity of ≥83% [95%CI:0.761-0.897]. True Positive (TP) of all SNVs = 589; False-Negative (FN) of all SNVs = 0. TPs of all indels = 105; FNs of all indels = 20. The undetected indels were manually checked on IGV. We observed that 12 of 20 were located +4 bp upstream of the exon (our bed file covers ±5 bp), four had poor coverage, two fell in highly repetitive regions and one was a long indel (18bp).

To determine specificity, we used the cancer-specific data from the four cell blends harbouring a total of 61 true positive and 87 true negative SNVs (Supplemental Table S7B). There were insufficient true negatives (n=3) to determine specificity for indels. SNVs were called and their presence was compared to the list of variants expected in the capture regions from the cell blends. The specificity of cancer-specific SNVs was ≥98% [95%CI:0.946-1]. Positive-Predictive Value (PPV) was ≥98% [95%CI:0.926-1] and the Negative-Predictive Value (NPV) was ≥98% [95%CI:0.946-1].

The range of VAF for the SNVs detected by our pipeline, including the background and the cancer specific variants (528 + 61 = 589), was 23% at ≥ 50% VAF (134/589), 35% at 50-20% of VAF (207/589) and 42% at < 20% (248/589). The range of VAF for the indels detected by our pipeline including the background and cancer specific indels (90 + 15 = 105) was 0% at > 50% of VAF (0/105), 31% at 50-20% of VAF (33/105) and 69% <20% (72/105).

### Performance and Variant Detection comparison in paired FF-FFPE clinical samples

To assess the performance of the panel on real clinical material we compared 15 paired clinical DNA samples isolated from both FF and FFPE samples. For the FFPE samples, we obtained an average of 93.4% ± 5.42% and 80.3% ± 20.3% of targeted positions covered at depths of ≥ 100x and ≥ 250x respectively. The overall mean depth for FFPE was 785 ± 333. Overall percentage of bases from unique reads on target for FFPE was 47.6% ± 2.3%. For FF samples, we obtained an average of 96.6% ± 0.6% and 90.9% ± 1.9% of targeted positions covered at depths of ≥ 100x and ≥ 250x respectively. The overall mean depth for FF was 977 ± 142. Overall percentage of bases from unique reads on target for FF was 44% ± 2.2%. As expected, duplicates were substantially lower in FF samples (54.5% for FFPE vs 29.9% for FF). Insert size for the library pre-capture DNA was 285 bp ± 24 for FFPE and 326 bp ± 24 for FF (Table 4).

**Table 4.** Comparison of quality metrics between formalin-fixed paraffin embedded (FFPE) and fresh frozen (FF) matched samples. Data expressed as means ± standard deviation.

|  | *Total reads* | *Percentage of reads mapped* | *Percentage of duplicates* | *Percentage of unique on target* | *Mean depth* |
| --- | --- | --- | --- | --- | --- |
| **FFPE** | $8.7 \times 10^7 \pm 3.4 \times 10^6$ | 95.5±2.2 | 54.5.2±9.2 | 47.6±2.3 | 785±333 |
| **FF** | $7.7 \times 10^6 \pm 1.2 \times 10^6$ | 98.6±0.7 | 29.9±6.9 | 44.2±2.2 | 977±142 |

VAFs found in the paired FF-FFPE samples were compared, obtaining an overall correlation of r^2^ = 0.983 (95%CI: 0.984-0.985; p<0.0001) (Figure 3 and Supplemental Figure S5). A total of 42.3% (5562/13146) variants were detected in FF but not in FFPE, of which 78.1% (4346/5562) had VAF below 5%, with 17.6% (982/5562) having VAF between 5-10%. Less than 5% variants missed in FFPE samples were present in FF at VAF above 10%.

**Figure 3.**
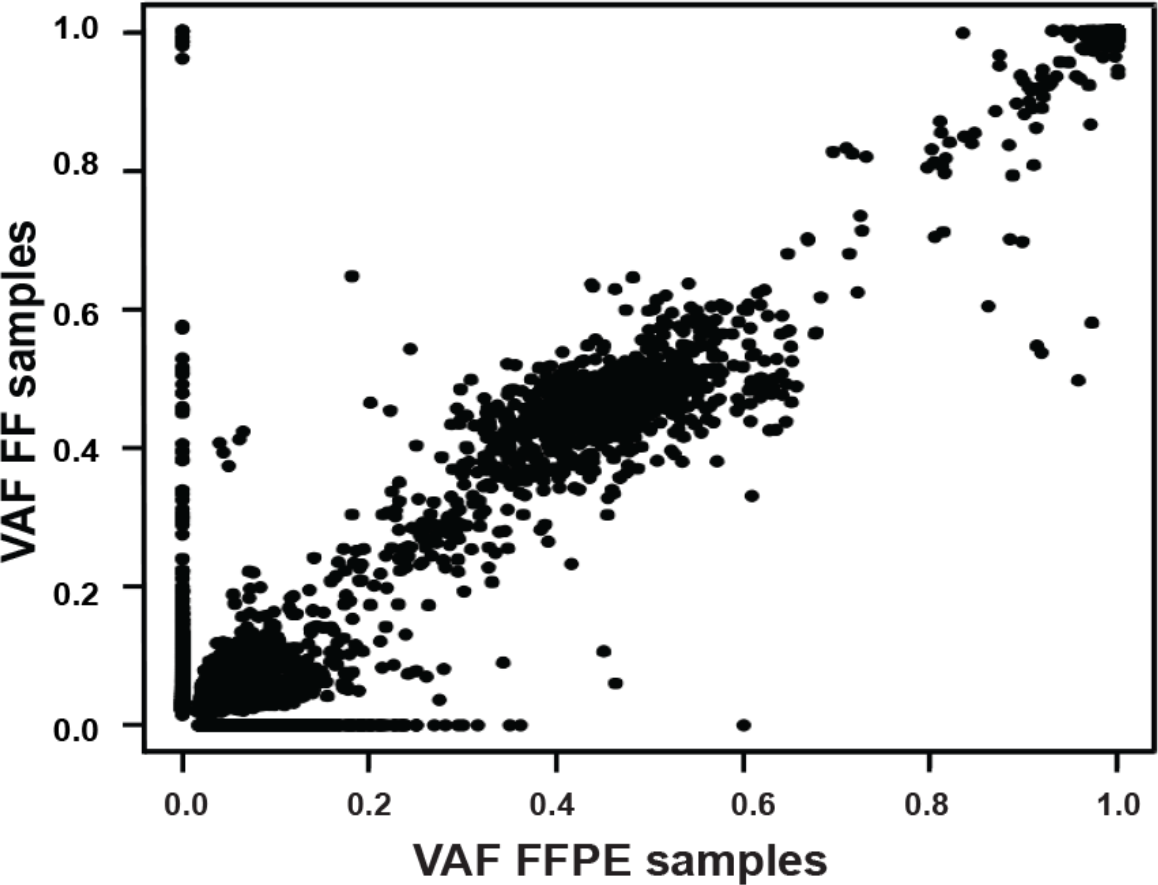
Overall correlation of variant allele frequency (VAFs) found between the 15 formalin-fixed paraffin embedded (x axis) and fresh frozen (y axis) paired samples.

Conversely, a total of 8.2% (1084/13146) variants were detected in FFPE but not in FF, of which 50.8% (551/1084) had VAF below 5%, with 33.2% (360/1084) having VAF between 5-10%, and the remaining 16.0% (173/1084) were present in FFPE only at VAF above 10%.

### Detection of known variants in paediatric samples

To assess the ability of the panel to detect known variants in clinical samples, we performed a variant analysis of 41 paediatric samples with 90 known genetic abnormalities (30 alterations in 13 cell lines and 60 alterations in 14 FFPE and 14 FF samples with known genetic alterations identified by routine testing): 50 SNVs, including mutations in *TP53, ALK, CTNNB1, DDX3X, SMARCA4*, one duplication (*BRAF* p.Thr599dup), 7 indels including *DDX3X and TP53*, 13 amplifications including *MYCN* and *CDK4*, and 19 chromosome/gene losses, for example chr 9q loss including loss of *PTCH1* and *TSC1*. 100% of the variants interrogated by the panel were successfully detected (Tables 5 and 6 and Supplemental Table S9).

**Table 5.** Known variants in paediatric cancer cell lines were compared against capture sequencing from the Cancer Cell Line Encyclopaedia and other published data.

| Cell line ID | Gene | Alteration | Detected | Allele frequency expected | Allele frequency observed |
| --- | --- | --- | --- | --- | --- |
| Be(2)C | *TP53* | p.C135F | YES | no data available | 100% |
| Be(2)C | *MYCN* | AMPLIFICATION | YES | no applicable | no applicable |
| CCA | *KRAS* | p.Q61L | YES | no data available | 29% |
| IMR32 | *ATM* | p.V2716A | YES | 59% | 59% |
| IMR32 | *MYCN* | AMPLIFICATION | YES | no applicable | no applicable |
| KELLY | *ALK* | p.F1174L | YES | 39% | 32% |
| KELLY | *MAP2K1* | p.A390T | YES | 48% | 47% |
| KELLY | *TP53* | p.P177T | YES | 93% | 99% |
| KELLY | *MYCN* | AMPLIFICATION | YES | no applicable | no applicable |
| LAN1 | *ALK* | p.F1174L | YES | no data available | 47% |
| LAN1 | *TP53* | p.C182* | YES | no data available | 99% |
| LAN1 | *MYCN* | AMPLIFICATION | YES | no applicable | no applicable |
| LAN5 | *ALK* | p.R1275Q | YES | no data available | 50% |
| LAN5 | *MYCN* | AMPLIFICATION | YES | no applicable | no applicable |
| NBLS | *NF1* | splice_acceptor_variant c.6705-1G>T | YES | no data available | 42% |
| RD | *ATM\** | p.D273N | YES | 17% | 2% |
| RD | *NF1* | p.E977* | YES | 56% | 59% |
| RD | *NRAS* | p.Q61H | YES | 68% | 61% |
| RD | *TP53* | p.R248W | YES | 100% | 100% |
| RH30 | *CDK4* | AMPLIFICATION | YES | no applicable | no applicable |
| RH41 | *APC* | p.M526L | YES | 60% | 59% |
| RH41 | *TP53* | p.P152fs | YES | 100% | 100% |
| RMS559 | *FGFR4* | p.V582L | YES | no data available | 76% |
| SKNAS | *NRAS* | p.Q61L | YES | 45% | 46% |
| SKNAS | *RB1* | p.L477P | YES | 47% | 31% |
| SKNAS | *TP53* | DEL exons 10,11 | YES | no applicable | no applicable |
| SKNSH | *NRAS* | p.Q61L | YES | 15% | 23% |
| SKNSH | *SMARCA4* | p.R973T | YES | 32% | 45% |
| SKNSH | *CHEK2* | p.T410fs | YES | 59% | 44% |
| SKNSH | *ALK* | p.F1174L | YES | no data available | 36% |

\*ATM mutation in this cell line is subclonal and variation in AF is expected with on-going passages

**Table 6.** Known variants in paediatric formalin-fixed paraffin embedded (FFPE, n=14) and fresh frozen (FF, n=14) samples were compared against other platforms such as RNA seq, 450k array, Sanger Sequencing and *FISH*.

| Genes with alterations detected by other methodologies | Alteration | Tumour Type | Total cases expected | % of cases detected |
| --- | --- | --- | --- | --- |
| *DDX3X* | *SNV and indel* | *Medulloblastoma* | 6 | 100 |
| *PTCH1* | *SNV and indel* | *Medulloblastoma* | 5 | 100 |
| *TP53* | *SNV and indel* | *Medulloblastoma* | 3 | 100 |
| *MYCN* | *SNV* | *Medulloblastoma* | 2 | 100 |
| *MYCN* | *Amplification* | *Neuroblastoma (n=3)*<br>*Medulloblastoma (n=4)* | 7 | 100 |
| *CTNNB1* | *SNV* | *Medulloblastoma* | 5 | 100 |
| *H3F3A* | *SNV* | *Glioma* | 3 | 100 |
| *SMARCA4* | *SNV* | *Medulloblastoma* | 3 | 100 |
| *BRAF* | *SNV* | *Glioma* | 2 | 100 |
| *ALK* | *SNV* | *Neuroblastoma* | 1 | 100 |
| *HIST1H3B* | *SNV* | *Glioma* | 1 | 100 |
| *AKT1* | *SNV* | *Medulloblastoma* | 1 | 100 |
| *ACVR1* | *SNV* | *Medulloblastoma* | 1 | 100 |
| *PIK3CA* | *SNV* | *Medulloblastoma* | 1 | 100 |
| *MLL2* | *SNV* | *Medulloblastoma* | 1 | 100 |
| *chr 9q - (PTCH1, TSC1)* | *loss* | *Medulloblastoma* | 5 | 100 |
| *chr 10- (PTEN, SUFU, FGFR2)* | *loss* | *Medulloblastoma* | 4 | 100 |
| *chr 6- (HIST1H3B, HIST1H3C, ROS1, ARID1B)* | *loss* | *Medulloblastoma* | 2 | 100 |
| *chr 9- (CDKN2A/B, PTHC1, TSC1)* | *loss* | *Medulloblastoma* | 2 | 100 |
| *chr12- (MLL2, CDK4)* | *loss* | *Medulloblastoma* | 1 | 100 |
| *ATM LOH* | *loss* | *Medulloblastoma* | 1 | 100 |
| *chr 3p- (CTNNB1, STED2)* | *loss* | *Medulloblastoma* | 1 | 100 |
| *chr17- (TP53, NF1, HER2, PPM1D)* | *loss* | *Medulloblastoma* | 1 | 100 |
| *chr17p- (NF1, TP53)* | *loss* | *Medulloblastoma* | 1 | 100 |
| **Total** |  |  | **60** |  |

### Detection of Rearrangements

Five sarcoma FFPE samples were included in the analysis where translocations had previously been detected by RT-qPCR involving *EWSR1*. Rearrangements in *EWSR1* were detected in four out of the five FFPE samples (80%) leading to fusion genes of *EWSR1* with partners *ATF1* (detected in two samples), *FLI1* and *CREB1* (Supplemental Figure S6). The fusion not detected was *EWSR1-NR4A3*. This is too small a sample to confirm validation of the panel for detection of translocations at this stage and further work is in progress.

## DISCUSSION

Targeted therapies are already the standard of care for several molecular subgroups of adult cancers. *EGFR* mutations or *ALK* rearrangements in lung cancer, *BRAF* V600E mutations in metastatic melanoma and breast cancer patients harbouring *HER2* amplifications are examples of therapeutic biomarkers routinely used in the adult population [36–38]. The implementation of personalised medicine in paediatric oncology has remained challenging partly due to the low incidence of childhood cancer, accessibility of drugs and regulatory hurdles [39]. Nevertheless, the understanding of genetics in childhood cancer over the last decade has improved thanks to large sequencing initiatives across the world [31–33]. The updated World Health Organization Classification (WHO) classification of brain tumours based on molecular features is a clear example of the huge impact of applying molecular profiling to guide diagnosis and treatment with the potential to improve outcomes in childhood cancers [40].

We have developed an NGS targeted sequencing based diagnostic test to accurately detect clinically relevant genomic alterations across 78 cancer genes in routine FFPE as well as FF paediatric samples. The overall performance of our assay was excellent; from the 901 regions captured only 24 (<3%) failed the quality control metrics mainly as a result of being located in GC-rich regions, and should be noted for future panel design. VAF for known SNVs and indels were very similar in within-run and between-run replicates, demonstrating that the assay is repeatable and reproducible. SNVs were detected at a wide range of VAFs simulating the heterogeneity expected in cancer samples including 33 SNVs with an expected VAF of 4-5%. The detection of variants at low VAF is crucial, especially in samples with a low neoplastic cell content. Sensitivity was ≥98% for SNVs and ≥83% for indels and specificity ≥98% for SNVs. False-negative calls were mostly at low VAF (≤5%) and predominantly occurred at splice sites. Variants were analysed in exons and the surrounding 5 bp, but were not called by our pipeline if they occurred in the last base. This could be solved expanding the sequence covered by bed file at intron:exon boundaries, but the relevance of these variants remains unclear. There is currently no consensus as to the most appropriate minimum region of interest to cover at splice sites for clinical reporting and in many cases the biological meaning of these mutations are unknown. The latest guidelines recommend calling likely disrupted gene function in nonsense, frameshift, canonical ±1 or ±2 splice sites, initiation codon, and single exon or multi-exon deletion, all of which would be covered with our current pipeline [41].

We also compared the performance of paired FFPE-FF specimens obtaining comparable quality metrics between both tissue types, as well as a high overall correlation of VAF. This is particularly important as most clinical samples routinely available are derived from FFPE tissue where nucleic acid quality is generally compromised and chemically challenged, leading to DNA degradation and potential deamination or oxidation artefacts. The discrepancies of the variants observed between FFPE and FF were mainly at low VAF, below or at the lower limit of detection of our approach. The discrepancies of the variants above 10% could be explained by variation in neoplastic cell content between FF and FFPE and intra-tumour heterogeneity leading to sub-clonal alterations. Three of the samples with more striking differences were brain tumours which are well known as highly heterogeneous tumours [42, 43].

We verified the accuracy of our NGS approach in cell lines and clinical specimens (FFPE and FF) containing known genetic abnormalities previously characterized by other methodologies and obtained a high concordance (r^2^ = 0.983). The FFPE and FF samples used for the validation were a cohort of specimens from several hospitals across the world. We obtained reproducible and accurate results from different quality samples processed in different pathology laboratories, demonstrating the value of this approach for the development of national and international clinical trials in paediatric oncology.

Our data shows that this NGS approach can detect structural variants, including amplifications, deletions and chromosomal rearrangements. These types of variants are not generally detected with commercial amplicon-based NGS panels, despite being of critical importance for the clinical management and diagnosis of paediatric patients (e.g. *MYCN* amplification in neuroblastoma, *EWSR1* in Ewing’s sarcoma). Only one out of five chromosomal rearrangements involving *EWSR1* was not identified by the assay which could be due to the lack of coverage at the intronic genomic location of the breakpoint. As expected, this is one of the limitations of the methodology, as capturing intronic regions commonly involved in translocations poses challenges associated to the presence of repetitive sequence elements. This can be partially overcome by including capture baits for the breakpoint regions of the most common partner genes involved in the translocations.

In summary, we have developed a robust clinical test that can detect SNVs, small indels, copy number variation and with high reproducibility and repeatability in routine clinical FFPE samples from a variety of centres. Our approach has been incorporated into a pilot molecular profiling study for paediatric patients at the Royal Marsden Hospital (London, UK) and this has now been extended across the UK as the METEOR programme, an interim step towards the UK’s more advanced paediatric molecular profiling programme, Stratified Medicine-Paediatrics (SM-Paeds) which is about to be rolled out throughout the UK. The NGS panel will form a key part of the SM-Paeds programme, which is underpinning UK patient eligibility screening for several clinical trials including the highly innovative international ITCC basket trial, called ESMART (NCT02813135), where patients are enrolled according to molecular alterations found in their tumours on biopsy at relapse. This is the first time that genomic results are incorporated into the patient’s record in paediatric cancer in the UK within a clinically relevant timeframe of 3-5 weeks. Our data shows that this NGS assay can be an accurate and a practical platform for molecular stratification and identification of actionable targets required to accelerate personalised medicine clinical trials in childhood solid tumours.

## Methods

### Validation samples

A representative selection of common, poor risk paediatric tumours was used for the validation comprising 132 samples: i) Four cell blends with validated variants (Tru-Q1-4 HorizonDiscovery, Cambridge, UK), ii) 15 paediatric cell lines iii) 83 FFPE clinical samples and iv) 30 FF clinical samples (Supplemental Table S3).

Local institutional review board approval was obtained for the project in addition to separate approvals from the contributing tumour banks (The Children’s Cancer and Leukaemia Group Tumour Bank and the Queensland Children’s Tumour Bank).

### Sample preparation

Assessment from haematoxylin and eosin (H&E) stained slides was performed by experienced pathologists to mark the region of the section containing tumour and to estimate neoplastic cell content, defined as the percentage of neoplastic cells out of total nucleated cells in the marked area. Tumour cellularity, reflecting the density of tumour nuclei, was also estimated. Macro-dissection of the marked area was performed, when appropriate, to enrich the tumour content. DNA from blood and cell lines, FF and FFPE samples was extracted using the QIAamp DNA blood mini kit, the QIAamp DNA mini kit and the QIAamp DNA FFPE tissue kit (Qiagen, Hilden, Germany), respectively. For specimens where DNA was extracted at local centres, methods are provided in supplementary methods. DNA was quantified using Qubit dsDNA High Sensitivity Assay Kit with the Qubit 2.0 fluorometer, (Invitrogen, Carlsbad, CA). Analysis by TapeStation 2200 using the genomic DNA ScreenTape assay (Agilent Technologies, Santa Clara, CA) was performed to determine the degree of fragmentation of genomic DNA prior to library preparation. Based on optimization studies, samples yielding DNA with median fragment length > 1000 bp were processed using 200 ng DNA. Samples with DNA < 1000 bp were processed using 400 ng if there was sufficient DNA.

### Gene Panel Capture and Sequencing

Library preparation was performed using the KAPA Hyper and HyperPlus Kit (Kapa Biosystems, Wilmington, MA, USA) and SeqCap EZ adapters (Roche, NimbleGen, Madison WI, USA), following the manufacturer’s protocol, including dual-SPRI size selection of the libraries (250-450 bp). In samples prepared using the KAPA Hyper Kit (n=39), DNA was sheared with the Covaris M220 (Covaris, Woburn, MA) using supplier protocols. KAPA HyperPlus employs enzymatic fragmentation and was used in 93 samples. Optimization of the process indicated that the change from enzymatic fragmentation resulted in a substantial improvement in library complexity and unique coverage depth compared to sonication [44]. Following fragmentation DNA was end-repaired, A-tailed and indexed adapters ligated. To optimise enrichment and reduce off-target capture, pooled, multiplexed, amplified pre-capture libraries (6 to 10 cycles according to the DNA input) were hybridized twice overnight (up to 13 samples per hybridization, consecutive days) using 1 µg of the pooled library DNA to a custom design of DNA baits complementary to the genomic regions of interest (NimbleGen SeqCap EZ library, Roche, Madison, WI, USA). A 5 cycle PCR was performed between hybridizations to enrich the captured product. After hybridisation, unbound capture baits were washed away and the remaining hybridised DNA was PCR amplified (12 cycles). PCR products were purified using AMPure XP beads (Beckman Coulter, Danvers, MA, USA) and quantified using the KAPA Quantification q-PCR Kit (KAPA Biosystems, Wilmington, MA, USA). Sequencing was performed on a MiSeq (Illumina, San Diego, CA, USA) with 75 bp paired-end reads and v3 chemistry according to the manufacturer’s instructions. For samples where germline matched control was available (n=23), pools from tumour and control DNA libraries were multiplexed separately for hybridization and combined prior to sequencing at a ratio of 4:1, increasing the relative number of reads derived from tumour DNA.

### Data Analysis

Primary analysis was performed using MiSeq Reporter Software (v2.5.1; Illumina), generating nucleotide sequences and base quality scores in Fastq format. Resulting sequences were aligned against the human reference sequence build GRCh37/Hg19 to generate binary alignment (BAM) and variant call files (vcf). Secondary analysis was performed in-house using Molecular Diagnostics Information Management System to generate QC, variant annotation, data visualisation and a clinical report. In the Molecular Diagnostics Information Management System, reads were deduplicated using Picard (http://broadinstitute.github.io/picard/), and metrics generated for each panel region. Oncotator (v1.5.3.0) (https://portals.broadinstitute.org/oncotator) was used to annotate point mutations and indels using a minimum variant allele frequency (VAF) of 5% and a minimum number of 10 variant reads. Manta (https://github.com/Illumina/manta) was used for the detection of structural variants. Variants were annotated for gene names, nature of variant (e.g. missense), PolyPhen-2 predictions, and cancer-specific annotations from the variant databases including COSMIC, Tumorscape, and published MutSig results. Copy number variation (CNV) was assessed using the ratio of GC-normalized depth of region of interest (ROI) in tumour against GC-normalized read depth of ROI in either matched germline DNA (when available) or the male cell line G147A (Promega, Madison, WI USA). Any ratio below 0.65 fold was defined as a potential deletion whereas a ratio above 2.4 was flagged as a potential amplification. All potential mutations, structural variants and CNVs were visualised using Integrative Genomics Viewer (IGV) and two individuals were required to review the mutation report independently. Variant calls from samples with previously known SNVs and indels were checked manually on IGV.

### Cell blends

The four cell blends contained 163 SNVs and 34 indels common to all four blends (background variants) (Supplemental Table S2A). Additionally, there were 61 SNVs and 17 indels, cancer variants, which were unique between blends, introduced at known VAF, and verified by ddPCR (Supplemental Table S2B). The four cell blends were used to assess overall performance, repeatability, intermediate precision, sensitivity and limit of detection. Specificity was determined using 87 true negative SNV sites (wild type) where another blend harboured a mutation at the corresponding position. The cell blends were processed and sequenced in two different runs by two independent users.

### Overall Performance

Four cell blends and five FFPE samples were used to measure performance across the capture design. The log mean depth across the panel was compared to the log depth of each region captured for each gene. Regions were classified as underperforming if the depth was lower than 2 x SD of the mean based on log_2_ [log_2_(ROI)>mean(log_2_(ROI))-2xSD(log_2_(ROI))]. GC content and mappability scores were compared against each region captured by the panel. Quality and coverage metrics were calculated across all the samples including i) total reads, ii) percentage of reads mapped to the reference sequence, iii) percentage of duplicates, iv) percentage of bases from unique reads de-duplicated on target, v) mean depth of targeted positions and vi) percentage of targeted positions with ≥50x, ≥100x and ≥250x coverage.

### Limit of detection

To assess the limit of detection and determine a reliable cut off for the analysis we used the unique cancer-specific set of variants from the four cell blends introduced at range of VAFs from 4% to 30%, defined by ddPCR.

### Precision

***Repeatability*** (or within-run precision) was determined by comparing the cell blend background variant data across the 4 different samples in the same run for variant detection and VAF. Intra-run pairwise correlation was calculated for two runs where the cell blends were prepared and sequenced by different users generating two sets of repeatability data.

***Intermediate precision*** (or between-run precision) was determined by comparing the cell blend background variant data between two runs for variant detection and VAF. Between-run pairwise correlation was calculated from two different runs prepared by different users and sequenced on different MiSeq instruments.

### Sensitivity and specificity

The sensitivity of the panel was determined by separately comparing the cell blend background variants and the cancer-specific variants introduced at known VAF. Specificity was determined using the cell blend cancer-specific set of data with known variants and known true negative sites. Variants were classified according to the different ranges of frequencies of the variants present in the DNA blends. We also determined Positive-Predictive Value and Negative-Predictive Value.

### Correlation between NGS targeted panel and other methodologies

13 paediatric cancer cell lines were tested harbouring a total of 30 known SNVs, deletions and amplifications previously identified by the Cancer Cell Line Encyclopaedia using Target Enrichment Sequencing (Agilent Technologies, Santa Clara, CA) and other published data [45–50]. Furthermore 33 samples (FF=14, FFPE=19) had a total of 65 known genetic alterations including i) SNVs detected by Sanger Sequencing (*H3F3A, TP53, CTNNB1, HIST1H3B, ALK, BRAF*) [51–53] and RNA-Seq ii) copy number changes by FISH (*MYCN*) [54] and 450k array and iii) rearrangements by Real-Time Quantitative PCR involving *ESWR1* as previously described [55, 56] (Refer to Supplemental Methods).

### Fresh frozen vs FFPE samples

15 paired FF and FFPE paediatric samples were compared for quality control metrics, coverage and the distribution of library inserts sizes between FFPE and FF paired samples. In addition, we correlated the VAF of the total variants found in the paired samples.

## Abbreviations

FFPE (formalin-fixed paraffin embedded); FF (fresh frozen); SNVs (single nucleotide variants); Indels (insertion-deletions); DIPG (diffuse intrinsic pontine glioma); NGS (next-generation sequencing); kb (kilobase); bp (base pairs); ddPCR (droplet digital PCR); SD (standard deviation); CI (confidential interval); HMW (high molecular weight); VAF (variant allele frequency); TP (true positive); FN (false negative); PPV (positive-predictive Value); NPV (negative-predictive Value); WHO (world health organization); H&E (haematoxylin and eosin); BAM (binary alignment); vcf (variant call files); CNV (copy number variation); ROI (region of interest); IGV (integrative genomics viewer).

## Author contributions

E Izquierdo designed and performed experiments, analysed-interpreted data, and wrote the manuscript; L Yuan analysed-interpreted the data and constructed analytical and visualisation tools; S George, L Chesler helped with the design, provided samples and gave advice; P Proszek performed experiments; C Jones, J Shipley, SA Gatz, L Marshall, C Stinson, AS Moore, SC Clifford, D Hicks, J Lindsey, R Hill, TS Jacques and J Chalker provided samples and/or gave advice; A Pearson, L Moreno and D Gonzalez de Castro conceived and designed the study; B Walker and D Gonzalez de Castro, supervised the research, interpreted data and reviewed the manuscript; D Gonzalez de Castro, K Thway and SO Connor carried out histopathological analysis; L Moreno reviewed the manuscript; M Hubank reviewed and edited the manuscript. All authors read and approved the final manuscript.

## Acknowledgments

We are enormously grateful to the Christopher’s Smile charity (grant numbers CSM 002 and CSM 003) for their support and enthusiasm to provide children with more effective and less toxic targeted drugs through molecular profiling. We also thank the contribution of Children’s Cancer and Leukaemia Group (CCLG).

## Conflicts of interest

The authors declared no conflict of interest with the submitted paper.

## Financial Support

This work was supported by Christopher’s Smile charity (CSM 002 and CSM 003) and the the NIHR Biomedical Research Centre at the Royal Marsden and the Institute of Cancer Research in London (Grant ref. A113).

## Supplementary Tables

**Supplemental Table S1.** (A) List of the 78 genes cover by the paediatric panel. The genes were classified according to whether alterations in that gene have been shown to be predictive biomarker (level 1), prognostic biomarker (level 2), diagnostic biomarker (level 3), potentially targetable biomarkers with inhibitors available or under development (level 4), known germline or high risk single nucleotide polymorphism (level 5) or unclear significance, research only (level 6). The table include tumour type were alterations have been reported, molecules targeting those genes and clinical trials available. (B) List of the 901 target regions included in the capture panel including exonic and intronic positions selected per gene. *Please note that intron region chosen for PAX3/7 should have been intron between exons 7 and 8 (PAX3: NM_181457, PAX7: NM_002584) instead of intron between exons 6 and 7*.

**Supplemental Table S2.** (A) List of background single nucleotide variants (SNVs, n=163) and insertion-deletion (indels n=34) on the four HD cell blends. (B) List of cancer-specific variants at known variant allele frequency (SNVs, n=61; indels n=17) and wild type sties used to determine specificity (n=87) on the four HD cell blends.

**Supplemental Table S3.** List of all samples used, including: tumour content, cellularity, tumour type and known genetic alterations.

**Supplemental Table S4.** QC used to determine underperforming regions across the four HD cell blends and 4 formalin-fixed paraffin embedded samples (FFPE). (A) Mean coverage for each targeted region. (B) Underperforming targeted regions. (C) Percentage of GC content in each he targeted region.

**Supplemental Table S5.** Quality Control metrics for the 132 samples used.

**Supplemental Table S6.** Coverage metrics for the 132 samples used.

**Supplemental Table s7.** (A) Variant allele frequency observed by the panel against expected by droplet digital PCR on the HD cell blends for the cancer specific single nucleotide variants and insertion-deletions (SNVs, n=61 and indels n=17) and (B) List of the cancer specific true negative sites (SNVs, n=87 and indels n=3).

**Supplemental Table S8.** Variant allele frequency observed on the HD cell blends for the selected single nucleotide variants (SNV, n=132) and insertion deletions (indels, n=27) used for the assessment of precision, sensitivity and specificity.

**Supplemental Table S9.** Samples with known alterations by other methodologies including 13 cell lines with 30 alterations and 28 clinical samples (14 formalin-fixed paraffin embedded, FFPE and 14 fresh frozen, FF) with 60 alterations.

## Supplementary Figures

**Supplemental Figure S1.** Consistency of single nucleotide variants (SNVs) allele frequency in the four HD blends with identical background variants. (A). Run_user1, (B). Run_user2. In both runs pairwise correlation of 132 SNVs allele frequencies was r^2^≥0.994 [95%CI:0.9910.996].

**Supplemental Figure S2.** Consistency of indels allele frequency in the 4 HD blends with identical background variants. (A). Run_user1, (B). Run_user2. In both runs pairwise correlation of 22 indels allele frequencies was r^2^≥0.785 [95%CI:0.652-0.919].

**Supplemental Figure S3.** Pairwise correlation of variant allele frequency for each of the four HD blends between the two runs (x axis correspond to run_user1 and y axis to run_user2) (A). Single nucleotide variants (SNVs) and (B). Insertion-deletions (indels). Pair-wise correlation was r^2^≥0.995 [95%CI:0.993-0.997] for SNVs and r^2^≥0.827 [95%CI: 0.716-0.937] for indels.

**Supplemental Figure S4.** Overall correlation of variant allele frequency (VAF) for the HD blends between the two runs analysing the four samples together (x axis correspond to run_user1 and axis to run_user2) (A). Single nucleotide variants (SNVs) and (B). Insertion-deletions (indels). Overall correlation was r^2^≥0.996 [0.995-0.997] for SNVs and r^2^≥0.827 [95%CI: 0.716-0.937] for indels.

**Supplemental Figure S5.** Correlation of variant allele frequency found between the 15 formalin-fixed paraffin embedded (FFPE, x axis) and fresh frozen (FF, y axis) paired samples.

**Supplemental Figure S6.** *EWSR1-CREB1* translocation in a sarcoma sample known to be positive for this translocation (A) Integrative Genomics Viewer plot (IGV) identified by the panel and (B) electropherogram by Sanger sequencing.

